# Computationally designing antimicrobial peptides for *Acinectobacter Baumannii*

**DOI:** 10.1101/419820

**Authors:** Ayan Majumder, Malay Ranjan Biswal, Meher K. Prakash

## Abstract

*Acinetobacter Baumannii,* which is mostly contracted in hospital stays, has been developing resistance to all available antibiotics, including the last line of drugs, such as carbapenem. Because of its quick adaptation there is an immediate need to design new antibiotics, possibly antimicrobial peptides (AMPs) to which bacteria do not develop resistance easily. Our threefold goal was to curate the available activity of AMPs on the same strain of *A. Baumannii*, build a neural network model for predicting their activity and use it to rationally pre-screen for lead generation from the thousands of naturally occurring AMPs. By curating and analyzing the recent activity data from 81 AMPs on ATCC 19606 strain, we develop a quantitative AMP activity prediction model. We selected three other models with comparable performance against a test set with known activities. With the goal of inspiring further studies on AMP drug candidates and their rational shortlisting, we made activity predictions for the entire database of AMPs using all the models. To handle the uncertainty of training with a small data set, highlighted peptides which had consistent results from all models.

---

*Acinetobacter Baumannii* (*A. Baumannii*)^1^ mainly implicated in hospital infections and is responsible for 80% of the infections caused by *Acinetobacter* species. *A. Baumannii* can also be found on normal human skin, but it generally does not pose threat towards a healthy person,^2–5^ besides the not so frequent skin and soft tissue infection, infection in the surgical site, urinary tract infection etc.^6,7^ In the past 30 years, *A. Baumannii* has evolved into a multidrug resistant (MDR)^8–10^ opportunistic pathogen that selectively infects seriously ill patients in intensive care unit (ICU), trauma or burn patients. ^2–4,11^ The presence of intrinsic efflux pump and high rates of genetic adaptation, contributes to adaptation against the antibiotics.^12–14^ Besides, it also possesses several beta-lactamase genes which offer resistance against beta-lactam antibiotics. ^15,16^ *A. Baumannii* has also been developing resistance against carbapenem^17^ which had been one of the last line of drugs against it. Combination therapies such as of colistin, polymixin B, and tigecycline are used to treat MDR strains, but these are complex compared to a single drug when it comes to quantification of the effect and the validation of their safety.^18–20^ Due to the growing concern about MDR, there is a surge for new types of antimicrobial agents.

Antimicrobial peptides (AMP) are a fundamental part of the innate defense system and are reportedly present in organisms from bacteria and fungus to humans.^21,22^ Although several modes of AMP activity, including DNA damage,^23^ RNA damage^24^ and targeting a regulatory enzyme^25^ or proteins^26^ have been proposed, it is generally believed that the positively charged AMPs act by disrupting the bacterial membrane.^27–29^ Because of this fundamental difference in the mechanism compared to the traditional drugs, it is believed that the bacteria do not develop resistance easily against AMPs. ^21^ The low toxicity of AMPs towards human cells and their low tendency to result in resistant strains makes them an ideal rational choice as the next generation antimicrobial agents,^30–32^ possibly eventually becoming an effective drug for *A. Baumannii*.

Quantitative Structure and Activity Relationship (QSAR)^33^ is an approach in computer aided rational drug design, which uses biophysical or biochemical parameters of the molecules to develop a quantitative relation with the measured activities. Once validated, the computational model can be used for predicting the activities of the possible drug candidates and for pre-screening them. Recent studies have developed QSAR relation using 29 small molecule drug candidates which act on the oxphos metabolic path of *A. Baumannii*.^34^ As noted above, since with AMP based drugs, bacteria are less likely to develop resistance to, we focus on QSAR for AMPs against *A. Baumannii*.

Several experimental groups have independently evaluated the activity of AMP against *A. Baumannii*. The goal of the present work is threefold. We curated these experimental result against a single, well studied, target ATCC 19606 strain. We developed a computational model using neural networks to rationally predict the activity from the biochemical attributes of the AMP. Since *A. Baumannii* is a growing threat, while realizing the potential limitations of training on 81 peptides, we also predict the activity of all AMPs from the AMP database on the *A. Baumannii*.

The data on the activity of AMPs on *A. Baumannii* is scattered in literature. We curated the data mainly with the goal of developing a quantitative model, and hence restricted the focus to the most commonly studied strain. The activity data of AMPs on *A. Baumannii* ATCC 19606 strain was curated from different sources^35–47^ and is presented in **Supplementary table 1**. The curated AMP data set had the activity of 81 AMPs with length ranging from 10 to 43 amino acids and charges in the range +1 to +12. Of these, for 69 AMPs the MIC was available (referred to as exact MIC data), and for the remaining 12, only the lower bound of MIC (refered to as the lower bound data).

Overall, the comprehensive collection of the data on AMP activity allowed an analysis relative the various biophysical parameters (*P*_*i*_) which are commonly used for developing quantitative relation with activities: (1) charge, which draws the AMPs selectively to anionic membrane,(2) length, and how it is commensurate with the membrane thickness reflecting the activity^48,49^ (3) molecular weight, which gives an idea of the bulkiness and membrane penetration efficiency (4) hydrophobic moment (*µ*_*H*_),^50^ which quantifies the amphipathic characters required to form pores in the membrane, (5) aliphatic index,^51^ which indicates the aliphatic content of the peptide, (6) grand average of hydropathy (GRAVY) based on Kyte-Doolittle hydropathicity scale, ^52^ (7) *in vivo* aggregation propensity, calculated by using a web-based software AGGRESCAN^53^ and (8) *in vitro* (*β*-sheet) aggregation propensity, calculated by using TANGO software (with ionic strength 0.02 M, pH 7.0 and temperature 298 K).^54^ The *in vitro* aggregation, before interaction with the membrane can at times stop proteolytic degradation^55^ by the bacteria but in many other cases reduce the drug potency.^56,57^ Further, the aggregation propensity affects the Barrel-Stave^58^ and carpet mechanisms^48^ of action differently. The distribution of the eight parameters for all the curated AMPs are given in **Supplementary Figure 1** and their individual relation with MIC in **Figure 1**, which shows that each of the parameters individually is not sufficient to describe the activity.

**Figure 1:**
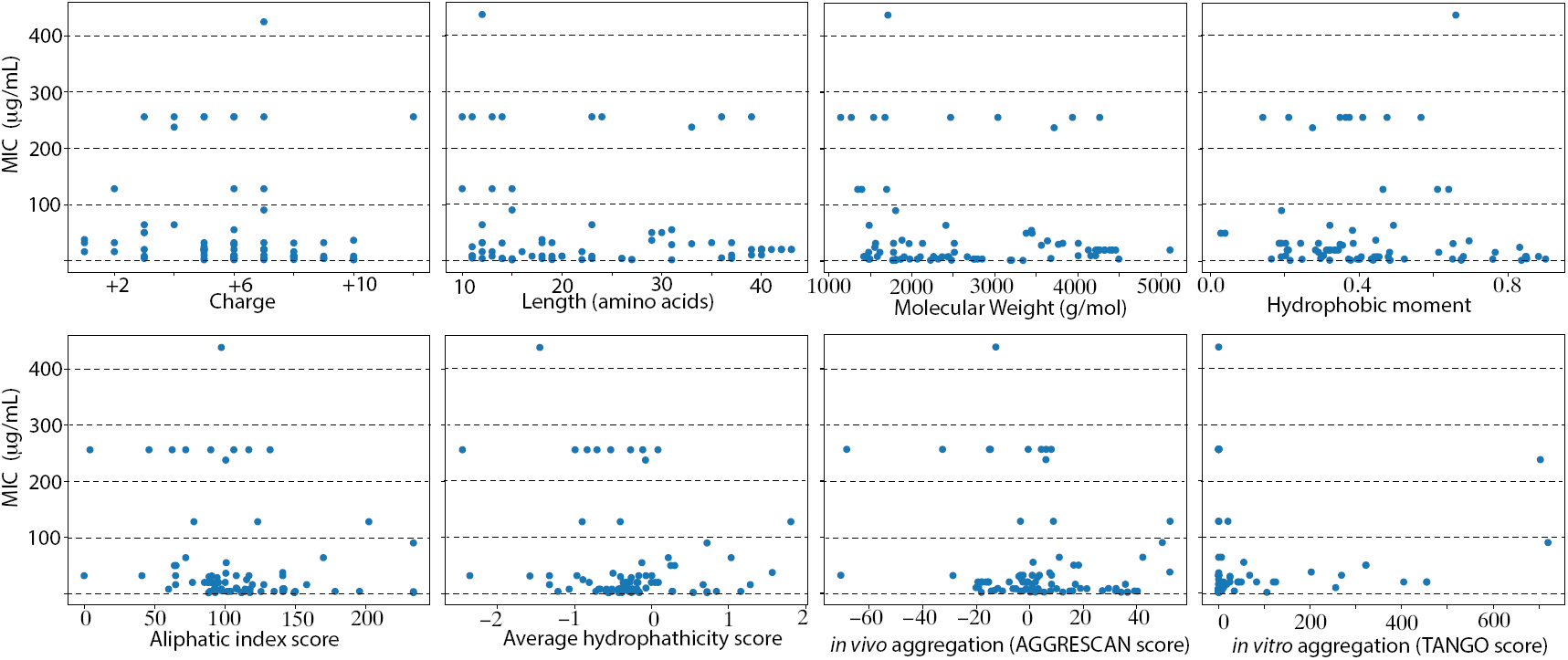
MIC versus different parameters. The AMP used in the analysis is shown in Supplementary Table 1.

Artificial neural network (ANN) method is used to obtain the relationship between the various above-mentioned parameters and the peptide activity (MIC values). Since the available data is limited, we did not discard the lower bound data in the quantitative model. We used 63 of the exact MIC values and 3 of the lower bound values, which were treated as the MIC for this analysis, for training (59 data points) and validation (7 data points). 6 data points with exact MIC were used for a quantitative test and the 9 remaining lower bound data points for an independent qualitative test. The activity of the AMPs was predicted by ANN model with an open module for machine learning called Scikit-learn^59^ in Python. For the activation function, logistic function was used and low memory BFGS optimization algorithm was used a solver. Two independent neural network calculations, one with a hidden layer of 6 neurons, another with 8 neurons were implemented for the quantitative activity prediction. 2500 trial runs in each case were made by taking 50 different random initializations for the input biases and 50 random choices for the training and validation sets. We screened the results of these 2500 trials with 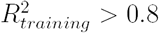 and 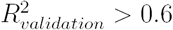, and identified 47 models with 6 neurons and 56 models with 8 neurons, to perform the test.

We created two independent test sets, one in which a quantitative MIC comparison was made and another quantitative one in which the calculated MIC was checked if it was more than the experimental lower bound. The selected models should simultaneously satisfy two conditions that at least 6 predictions in lower bound data set were correct and 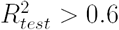 for the exact data. 4 models satisfied these conditions, with 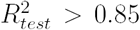 and they were selected. The best among these models (referred to as Model-1), had good predictions 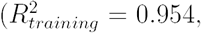, 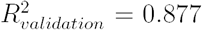 and 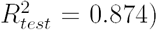. The experimental MIC for the quantitative data set versus MIC values predicted from Model-1 is shown in **Figure 2**. Results obtained from other three models are given in **Supplementary figure 2**.

**Figure 2:**
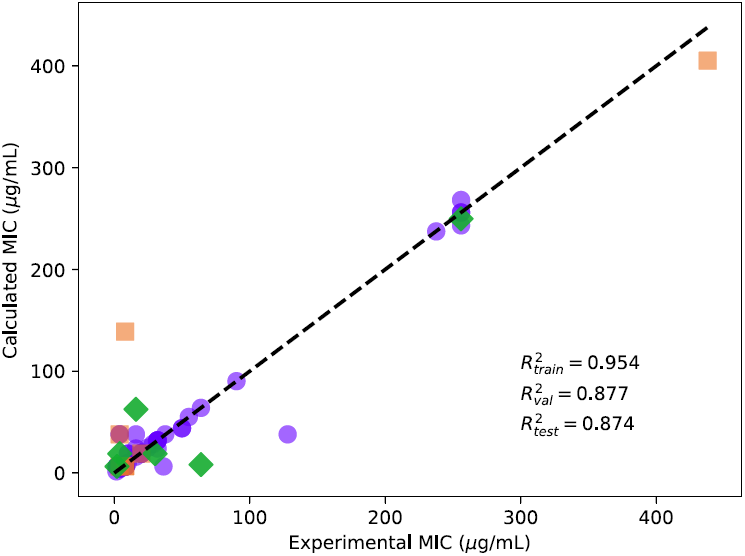
Comparison of the experimental and calculated MIC (*µ*g/mL) of curated AMPs on *A. Baumannii* obtained from Model-1. Training (purple circles), validation (orange squares) and test (green diamonds) sets are shown. The data used in the analysis is shown in Supplementary Table 1.

It is important to know, which are the parameters (*P*_*i*_) that are most responsible for the activity on *A. Baumannii*. In the combined training and validation set used for accepting the models, we replaced (*P*_*i*_) with its average *<P*_*i*_*>* and measure the difference 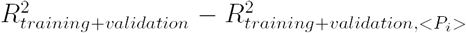. 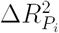 is treated as reflecting the importance of the parameter. The results obtained from Model-1 are given in **Figure 3** and those from the other three models are given in **Supplementary Figure 3**. From our calculations, we found out that the *in vitro* aggregation propensity in the most important parameter in aggregation propensity influences the outcomes of the predictions the most.

**Figure 3:**
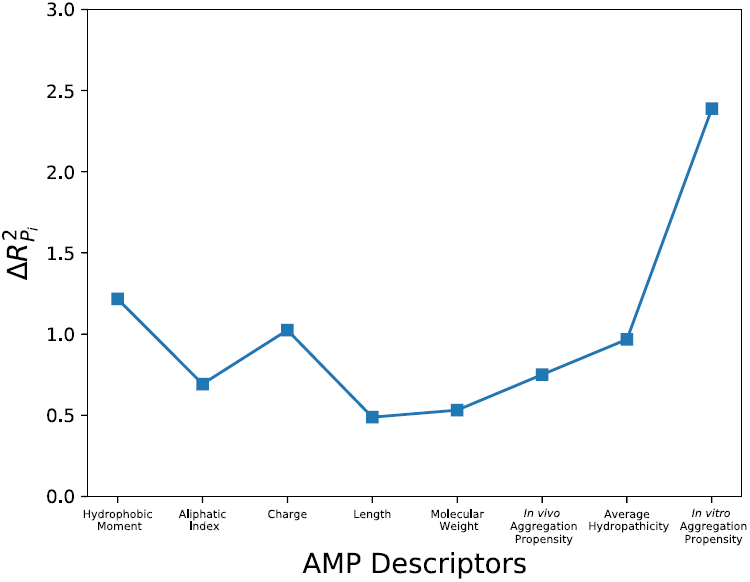
The relative importance of the different parameters in Model-1 is shown. *In vitro*

Considering the health threat *A. Baumannii* is posing, and the potential of AMPs for antibiotic-resistance-free activity, we propose a rational basis for the design of AMPs for *A. Baumannii*. Our models were used to predict the MIC values of the 2338 AMPs obtained from AMP database^60–62^ (https://aps.unmc.edu). In order to reduce the risk of a poorly trained ANN model with limited data, we filtered the predictions from all models for a consistent prediction that is within ΔMIC*≤* 5*µ*g/ml (**Supplementary table 3**).

To our knowledge ours is the only QSAR study for predicting AMP activity against *A. Baumannii*. The present work is different from the only the only other QSAR in two different ways, using AMPs in stead of small molecules for a better tolerance to antibiotic resistance and a slightly larger set (81 AMPs compared to 29 small molecules). Using the ANN models we developed, we could make quantitative predictions for the entire database of naturally occuring AMPs. We hope that our work will inspire the further studies quantifying the activity of AMPs on *A. Baumannii*, some of which may follow the activity predictions and others that differ offer an opportunity to retrain the ANN models.

## Supplementary Information

### Supplementary Table 1

**Supplementary Table 1:**
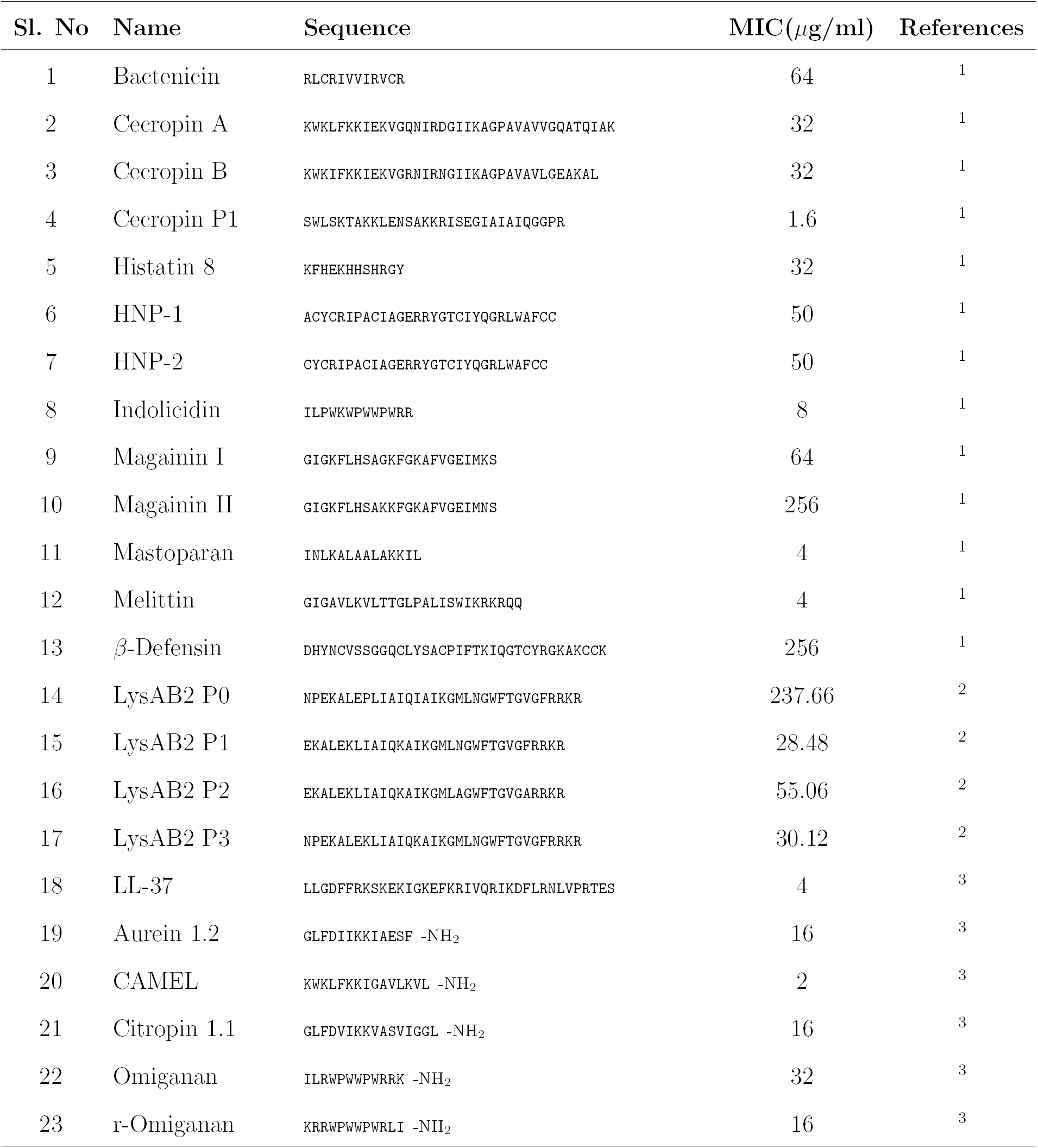
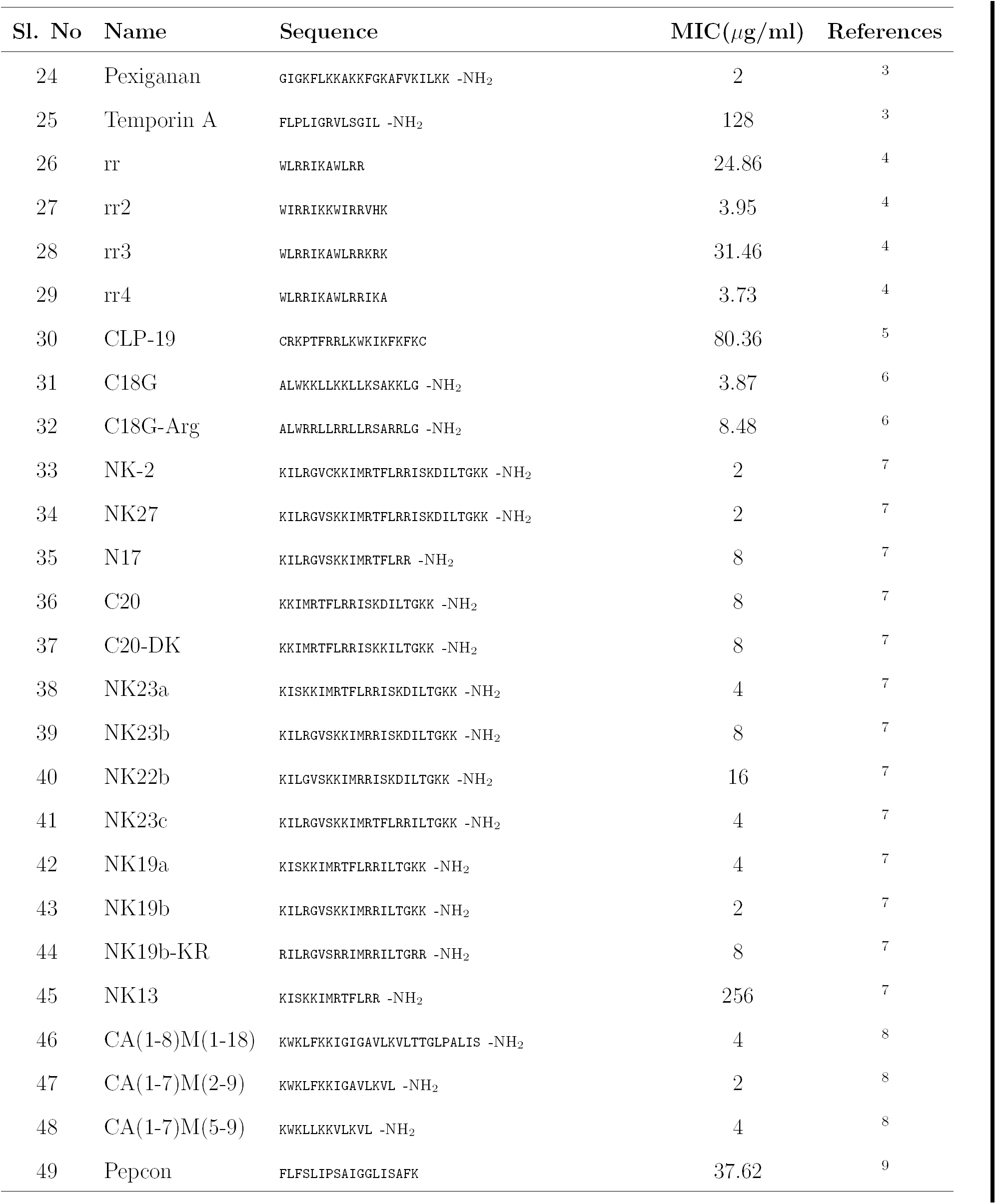
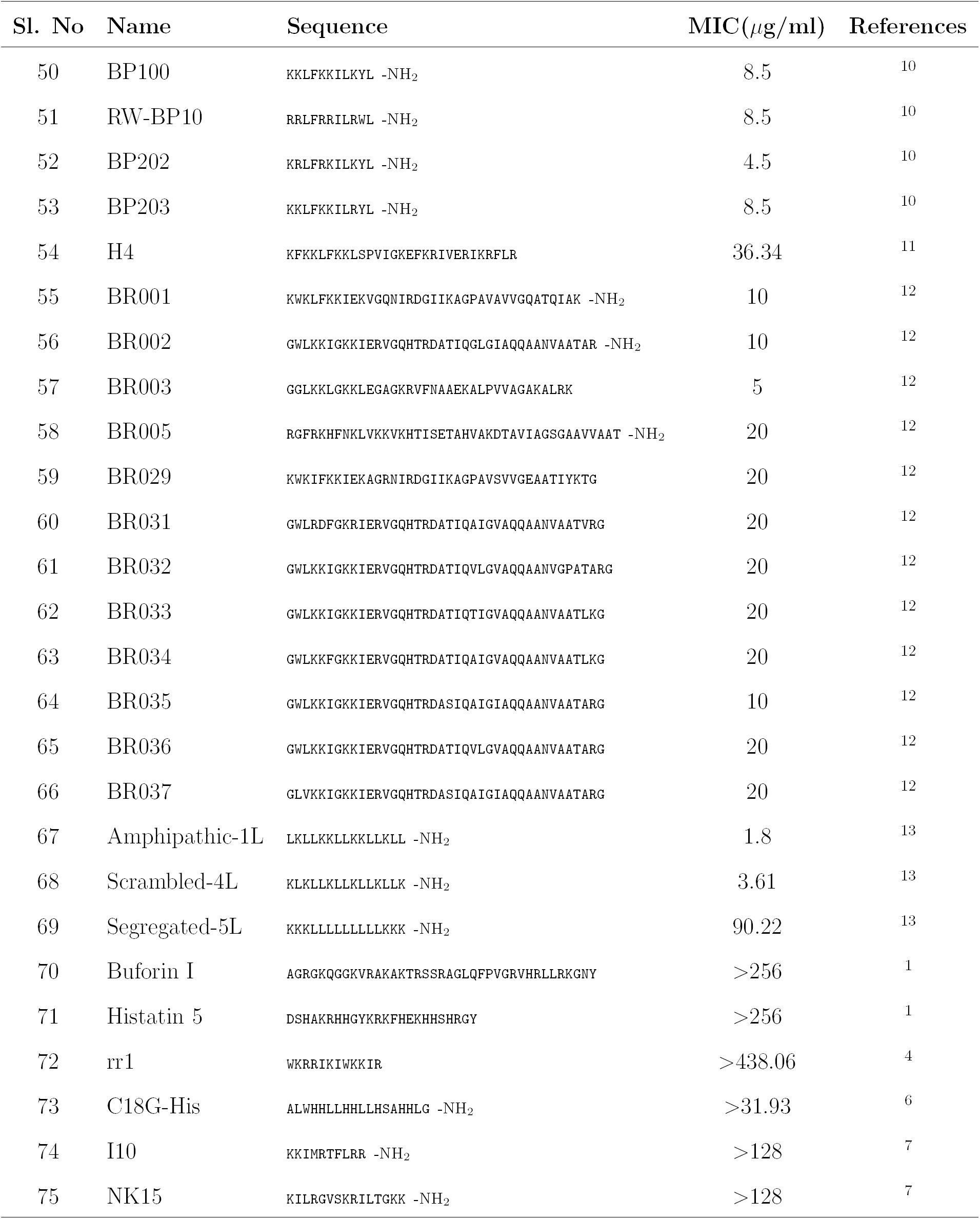
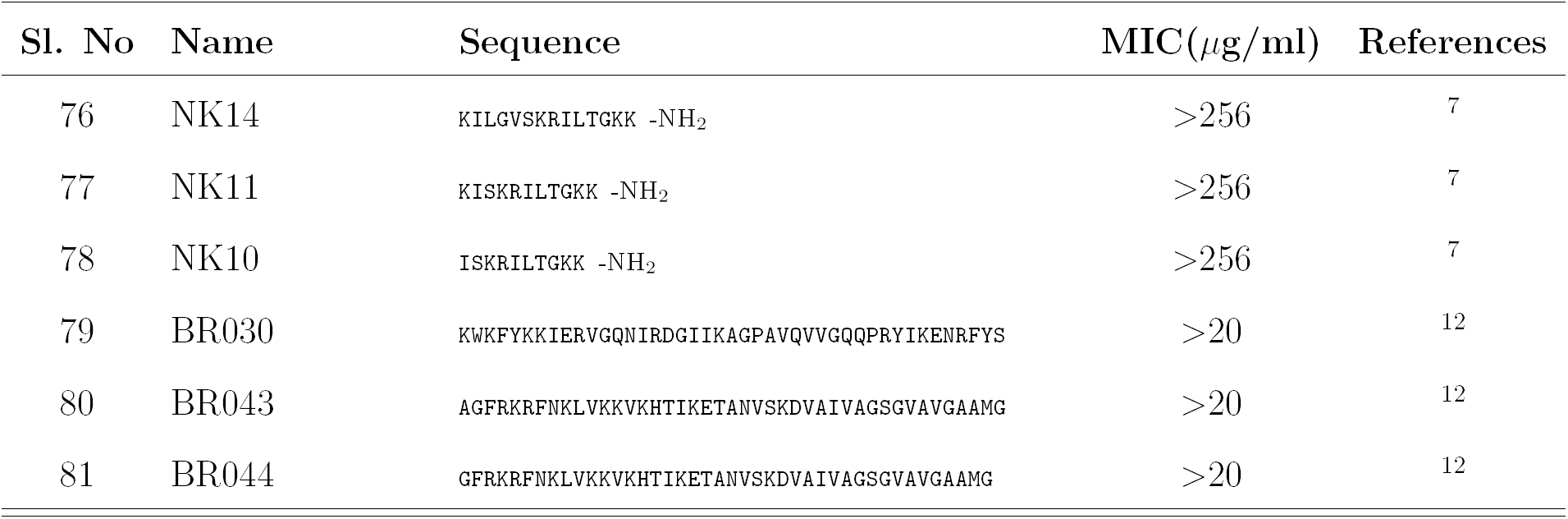
Details of all the 81 antimicrobial peptides used in the present work.

### Supplementary Table 2

**Supplementary Table 2:**
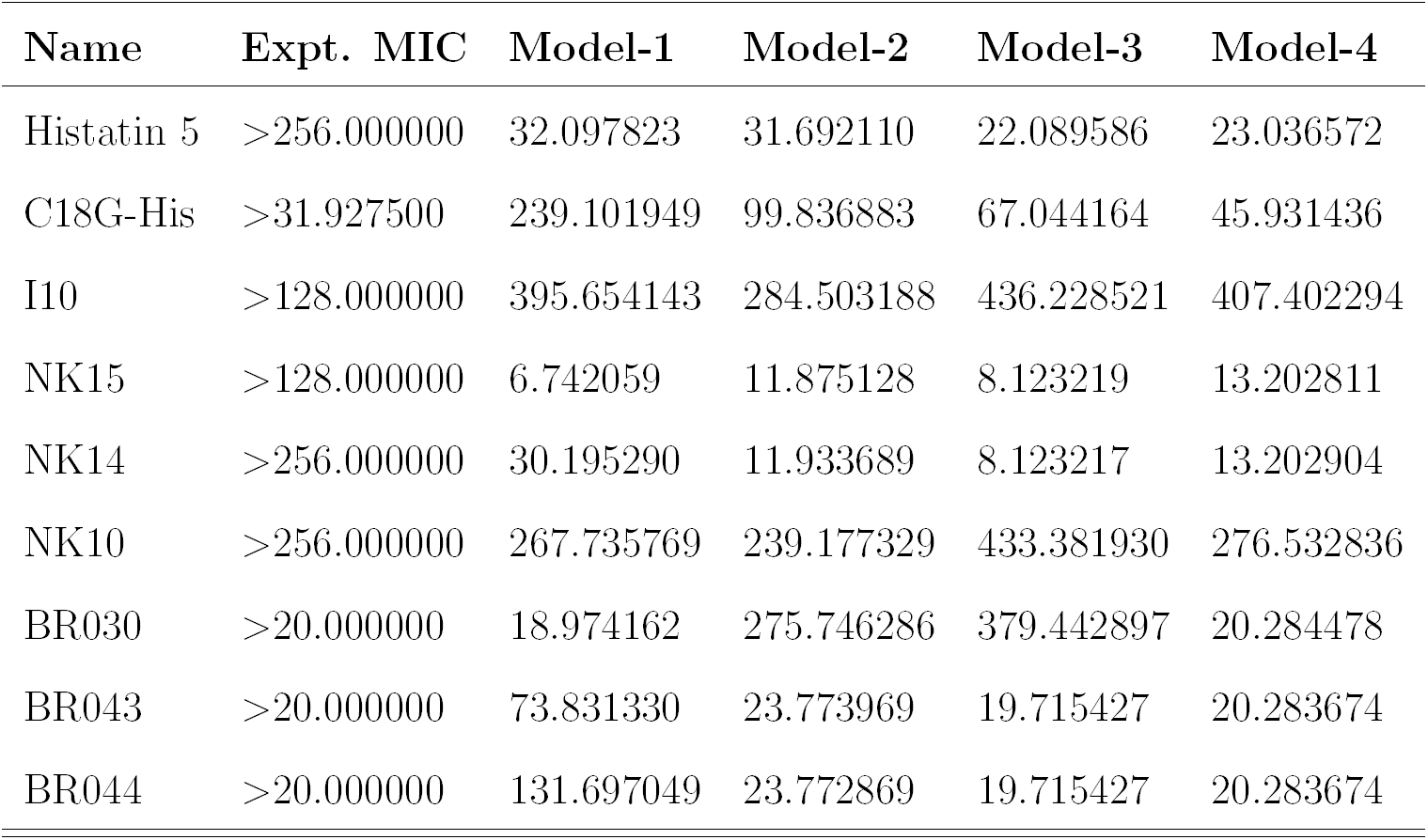
Details of the 12 lower-bound AMP data, these 9 were used for a qualitative test. Before accepting a model, at least 6 of these 9 results were verified to be satisfying the condition and then a quantitative test condition is used with the test data set from the exact data.

**Supplementary Figure 1:**
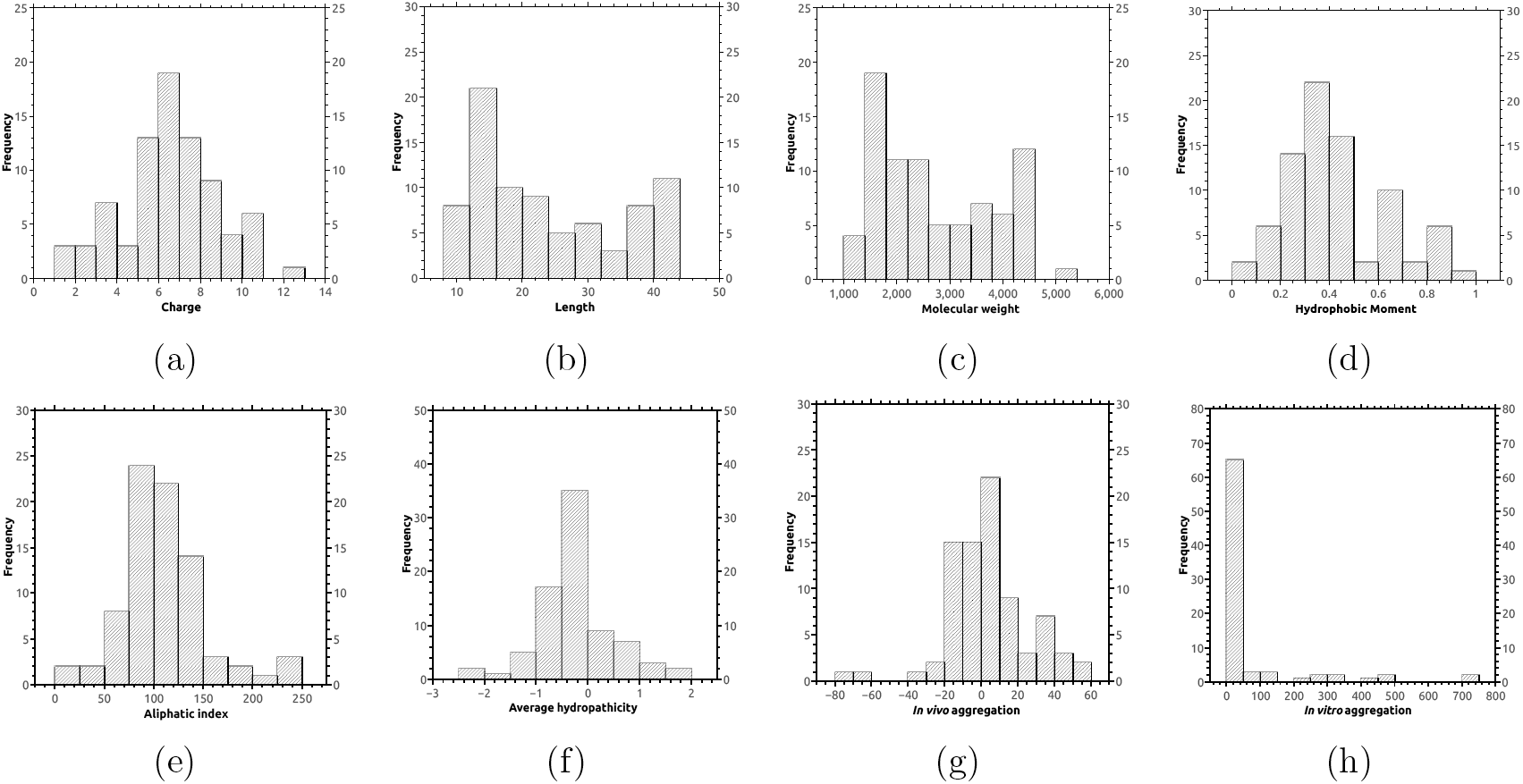
Histogram of all the parameters. The data used in the analysis is shown in Supplementary Table 1.

**Supplementary Figure 2:**
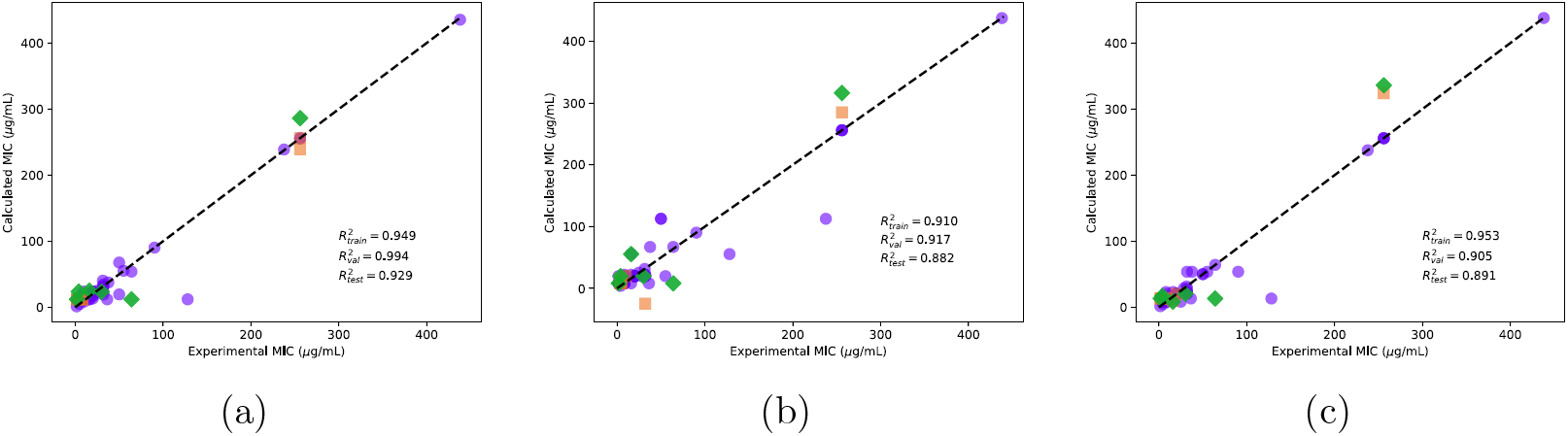
Comparison of the experimental and calculated MIC (*µ*g/mL) of curated AMPs on *A. Baumannii* obtained from (a) Model-2, (b) Model-3, (c) Model-4. Training (purple circles), validation (orange squares) and test (green diamonds) sets are shown. The data used in the analysis is shown in Supplementary Table 1.

**Supplementary Figure 3:**
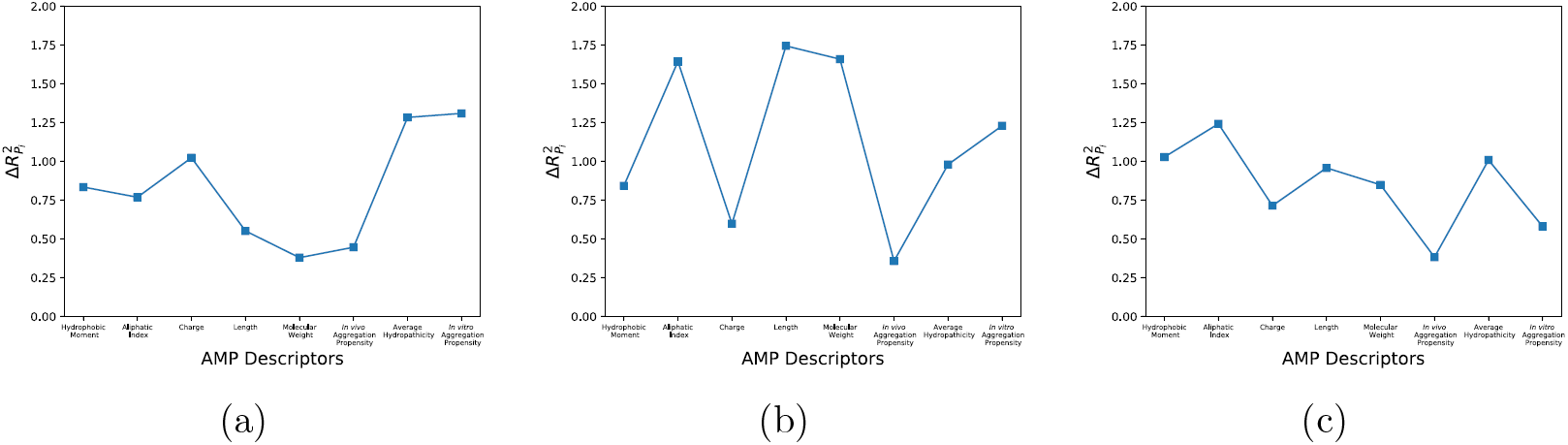
The relative importance of the different parameters is shown for Model-2, (b) Model-3, (c) Model-4.

### Supplementary Table 3

**Supplementary Table 3:**
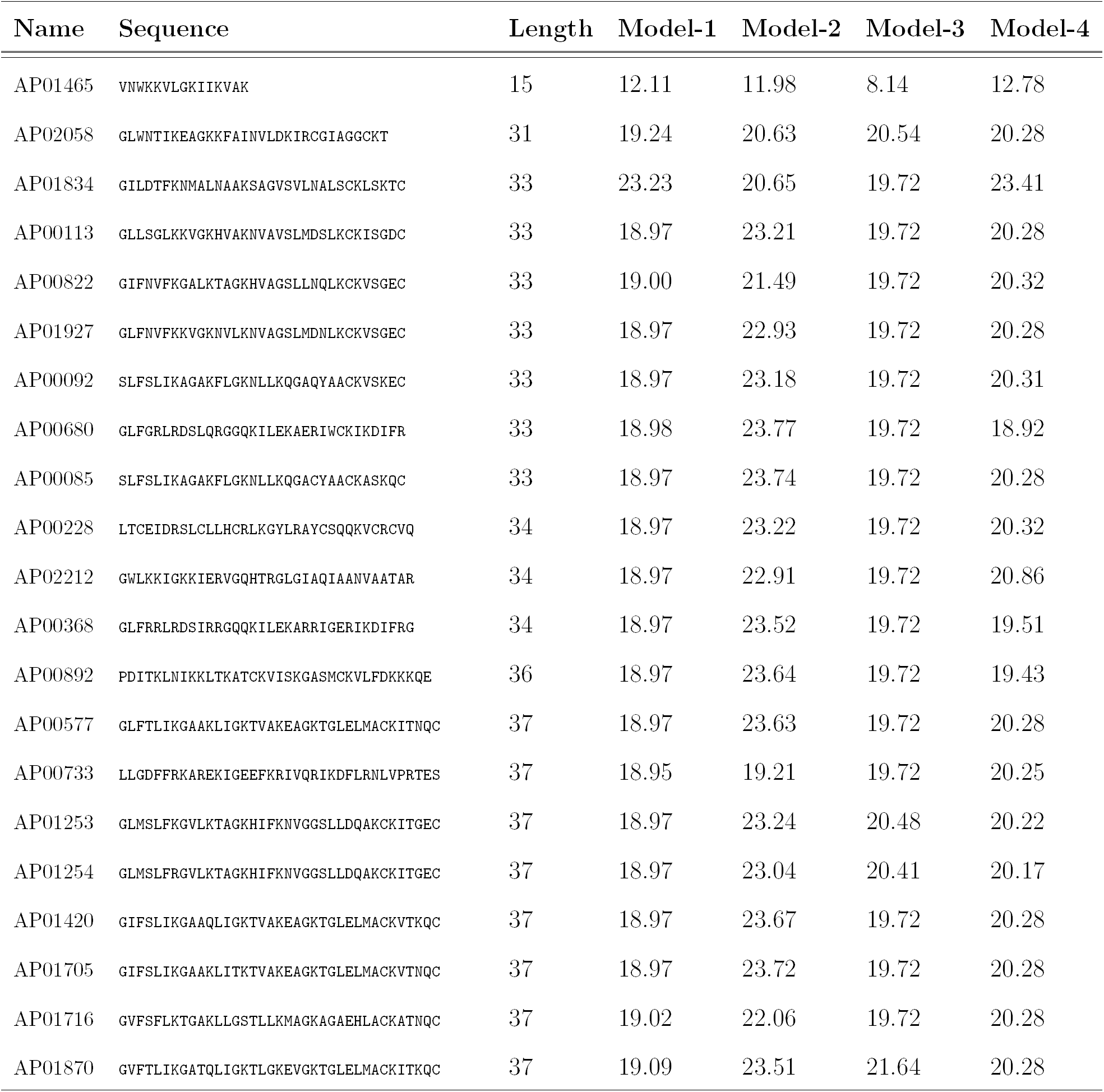
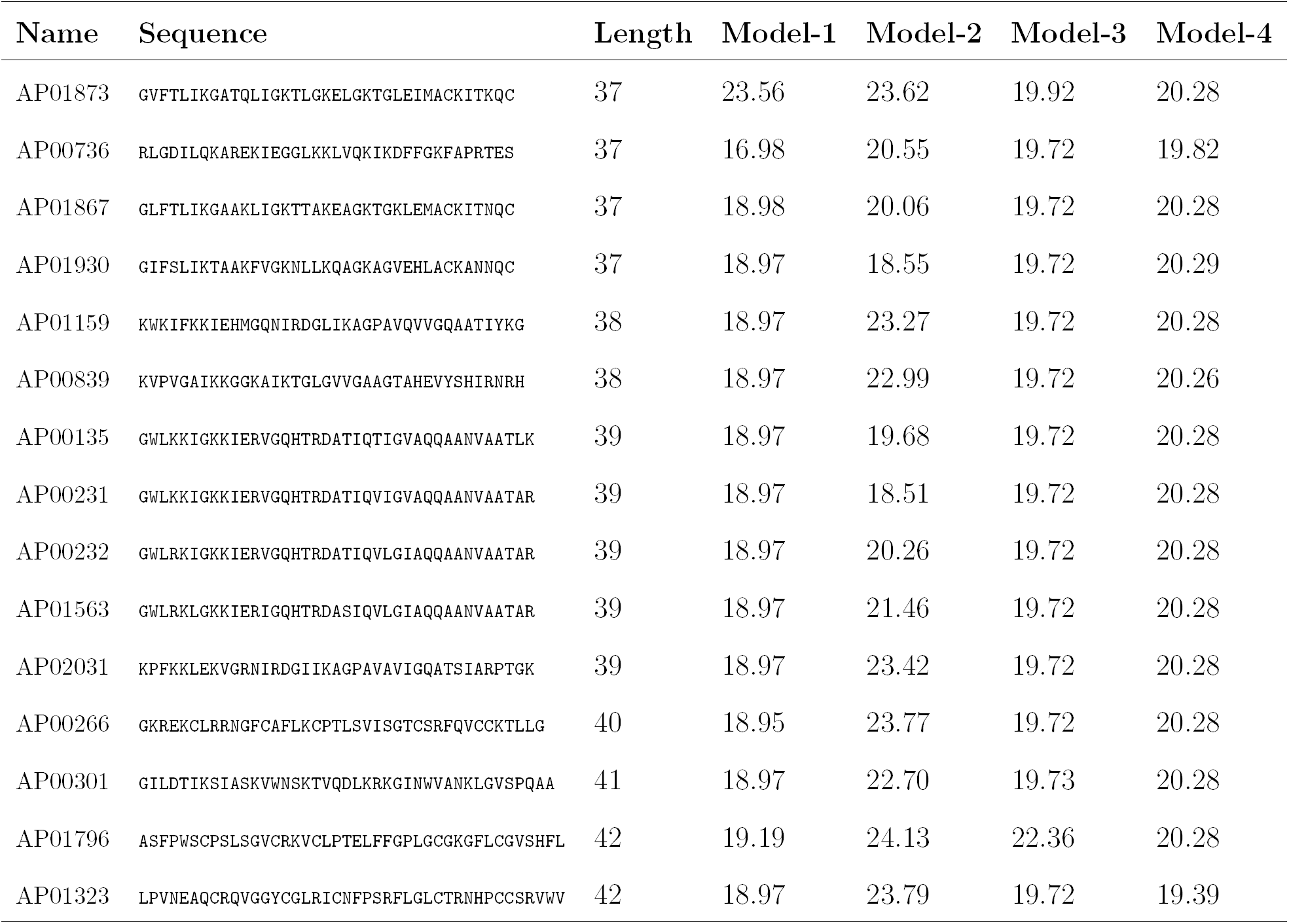
Using the 4 different models, we predicted the activity of 2338 naturally occurring AMPs documented in the AMP database. Of these the AMPs which had consistent predictions from any two models (ΔMIC ≤ 5 *µ*g/ml) were selected and presented in this table.

